# Reversing the miRNA -5p/-3p stoichiometry reveals physiological roles and targets of miR-140 miRNAs

**DOI:** 10.1101/2021.10.12.463698

**Authors:** Young Cameron, Caffrey Melissa, Christopher Janton, Tatsuya Kobayashi

## Abstract

The chondrocyte specific miR-140 miRNAs are necessary for normal endochondral bone growth in mice. miR-140 deficiency causes dwarfism and craniofacial deformity. However, the physiologically important targets of miR-140 miRNAs are still unclear. The miR-140 gene (*Mir140*) encodes three chondrocyte-specific microRNAs, miR-140-5p, derived from the 5’ strand of primary miR-140, and miR140-3p.1 and -3p.2, derived from the 3’ strand of primary miR-140. miR-140-3p miRNAs are ten times more abundant than miR-140-5p likely due to the non-preferential loading of miR-140-5p to Argonaute proteins. To differentiate the role of miR-140-5p and -3p miRNAs in endochondral bone development, two distinct mouse models, miR140-C>T, in which the first nucleotide of miR-140-5p was altered from cytosine to uridine, and miR140-CG, where the first two nucleotides of miR-140-3p were changed to cytosine and guanine, were created. These changes are expected to alter Argonaute protein loading preference of -5p and -3p to increase -5p loading and decrease -3p loading without changing the function of miR140-5p. These models presented a mild delay in epiphyseal development with delayed chondrocyte maturation. Using RNA-sequencing analysis of the two models, direct targets of miR140-5p, including *Wnt11*, were identified. Disruption of the predicted miR140-5p binding site in the 3’ untranslated region of *Wnt11* was shown to increase Wnt11 mRNA expression and caused a modest acceleration of epiphyseal development. These results show that the relative abundance of miRNA-5p and -3p can be altered by changing the first nucleotide of miRNAs in vivo, and this method can be useful to identify physiologically important miRNA targets.

## Introduction

Small regulatory microRNAs (miRNAs) suppress gene expression of target RNAs by directly binding to sequences mainly in the 3’-untranslated region (UTR) via partial base pairing (Bartel, 2018). The evolutionarily conserved and chondrocyte-specific microRNA gene, *Mir140*, is essential for normal endochondral bone development. Mice missing *Mir140* develop dwarfism, cranial deformity, and shortening of limbs (Nakamura et al. 2011; Miyaki et al. 2010; Papaioannou et al. 2013). In humans, we have recently reported that a single nucleotide substitution at the second nucleotide of miR-140-5p of *Mir140* causes a novel skeletal dysplasia, spondyloepiphyseal dysplasia (SED) MIR140 type, which is characterized by short stature, facial dysmorphism, and the unique epiphyseal deformity of long bones. Loss of *Mir140* in mice causes mild acceleration of chondrocyte differentiation (Nakamura et al. 2011) and a reduction in proliferation of growth plate chondrocytes (Miyaki et al. 2010), resulting in impaired longitudinal bone growth in the cranial base, spine, and limbs. However, despite the evidence that *Mir140* plays an important role in endochondral bone development and growth, molecular mechanisms by which *Mir140*-deficiency affects growth plate chondrocyte behavior are still largely unknown.

*Mir140* produces three miRNAs, miR-140-5p, derived from the 5’-strand of the primary miR-140 (pre-miR-140) miRNA, and miR-140-3p.1 and -3p.2, derived from the 3’-strand of pre-miR-140, after cleavage of the primary transcript by Drosha and subsequent cleavage of the loop region by the endonuclease, Dicer (Gebert and MacRae 2019; Suzuki et al. 2015). It is important to note that Dicer enzymatic cleavage occurs at two sites on the 3’-strand of the pre-miR-140 producing two species of miR-140-3p. Our previous study, investigating the regulatory effects of miR-140-5p and miR-140-3p on their predicted targets using *Mir140*-null chondrocytes, revealed that the loss of miR-140 miRNAs causes greater impacts on miR-140-5p target gene expression than miR-140-3p target gene expression despite the fact that miR-140-3p is substantially more abundant than miR-140-5p (Grigerioniene et al. 2019). These observations suggest that miR-140-5p has greater regulatory effects and that the miR-140-5p abundance might need to be suppressed to achieve optimal fitness for animals. From these findings, we hypothesize that reversing the miR-140-5p and miR-140-3p stoichiometry may elucidate the regulatory targets of miR-140-5p and miR-140-3p.

Upon generation of mature -5p and -3p miRNAs, miRNAs are loaded onto Argonaute (Ago) proteins where miRNA-target RNA interaction takes place (Gebert and MacRae 2019). Pre-miRNA processing by Dicer and Ago-loading is coupled, and therefore either the -5p or the - 3p strand is selected for Ago loading (Meijer et al. 2014). The Ago protein complex consists of four domains, MID, PIWI, N-terminal, and PAZ (Hutvagner and Simard 2008). The first monophosphorylated nucleotide of mature miRNAs binds to grooves of the MID and PIWI domains, and the region of the 3’-end of the miRNA associates with the PAZ domain (Sheu-Gruttadauria and MacRae 2017). Due to these physical association with Ago, miRNAs interact with target RNAs, mainly, via the second through the eighth nucleotides, called the “seed sequence”, which determines miRNA target specificity (Agarwal et al. 2015; Gebert and MacRae 2019). Strand selection for Ago loading appears to follow relatively simple thermodynamic rules (Suzuki et al. 2015). Upon Dicer cleavage, pre-miRNA is processed to a miRNA-5p and a -3p that form a double stranded RNA. It has been reported that the strand whose 5’ end is in the thermodynamically more unstable end of the miRNA duplex is preferred for subsequent Ago loading (Suzuki et al. 2015). In addition, the miRNA strand starting with 5’-uridine (U) or 5’-adenosine (A) is preferred for Ago loading (Suzuki et al. 2015). Based on the publicly available small RNA-sequencing data (Griffiths-Jones 2006) and our miRNA-seq data (Grigerioniene et al. 2019), although both miR-140-5p and -3p are expressed, the abundance of miR-140-3p is ~10 times greater than that of miR-140-5p. These findings are in accord with the aforementioned rule, since miR-140-5p starts with cytosine (C), a non-preferred nucleotide for Ago loading, and miR-140-3p miRNAs starts with U or A, preferred nucleotides for Ago loading. Consequently, we hypothesize that either changing the 5’ nucleotides of miR-140-3p miRNAs to less preferential nucleotides (C and G) and/or changing the 5’ nucleotide of miR-140-5p to a more preferential nucleotide (U) would reverse the relative abundance of miR-140-3p and miR-140-5p.

In this study, we present two mouse models in which the stoichiometric balance of miR-140-5p and -3p is reversed by altering the nucleotides of the 5’-end of miR-140-3p and miR-140-5p. These mutant mice present a mild delay in chondrocyte maturation. Using RNA-sequencing analysis of these two stoichiometric inverted mouse models, along with a previously reported miR-140 knock-out mouse model, we identified miR-140-5p targets, including *Wnt11*. Mice lacking the binding site for miR-140-5p in the *Wnt11* 3’ untranslated region showed increased *Wnt11* expression, and these mice presented a modest acceleration in epiphyseal development. These results provide evidence that miR-140-5p directly regulates *Wnt11* expression leading to regulation of endochondral bone development.

## Results

### Generation of mouse models with reversed miR-140-5p and -3p stoichiometry

In order to address the strand-specific miR-140 miRNA regulatory roles, two distinct miR-140 mutant mouse models were generated via CRISPR/Cas9-mediated gene editing. We hypothesized that changing the first nucleotide of miR-140-5p from C, a non-preferred nucleotide for Ago loading, into U, a preferred nucleotide for Ago loading, would increase the miR-140-5p abundance, and thus decrease miR-140-3p miRNAs. Additionally, we hypothesized that changing the first nucleotides of the 5’-end of miR-140-3p.1 and -3p.2 from U and A, preferred nucleotides for Ago loading, to C and G non-preferred nucleotides for Ago loading, would decrease miR-140-3p loading, and thus increase the abundance of miR-140-5p. Consequently, the first model, designated as miR-140-CG, was made by switching the first 5’ nucleotide of miR-140-3p.1 and -3p.2 U and A into C and G, respectively (Fig. 1A-B). In addition to the 5’ nucleotide switching in the miR-140-3p miRNAs, complementary mutations were made to the 3’ nucleotide end of miR-140-5p (Fig. 1A-B). The purposes for these secondary mutations were to maintain the hairpin-loop structure of the pre-miR-140 for normal processing by Drosha and Dicer and to increase the thermodynamic stability of the 5’ end of miR-140-3p of the miR-140-5p and -3p duplex upon cleavage. The thermodynamic stability of the 5’ end of -5p and -3p miRNA duplexes is another major determinant for strand selection where the miRNA strand in which the 5’ end is located in the thermodynamically unstable side is preferred for Ago loading (Schwarz et al. 2003; Khvorova et al. 2003; Suzuki et al., 2015). It is important to note that these changes are expected to have little or no impact on the targeting specificity of miR-140-5p because the 3’ end of miRNAs are associated with the PAZ domain of Ago proteins, and thus does not participate in the miRNA-target RNA interaction, while they should decrease Ago loading of miR-140-3p strands and increase miR-140-5p loading. As for the effect of these changes on miR-140-3p miRNAs, they do not affect the function of miR-140-3p.2 because its seed sequence is not altered, but they alter the first nucleotide of the seed sequence of miR-140-3p.1, and therefore disrupt its function. However, the hypothesis denotes that the expression of miR-140-3p miRNAs is expected to be significantly decreased, such that the functional change of miR-140-3p.1 should have a limited physiological impact. The second model, designated as miR-140-C>T, was created by mutating the first nucleotide of miR-140-5p from C to U (Fig. 1A-B). In this model, there is no seed sequence change in miR-140-5p or -3p miRNAs, and therefore, no functional changes of miR-140 miRNAs are expected.

**Figure 1.**
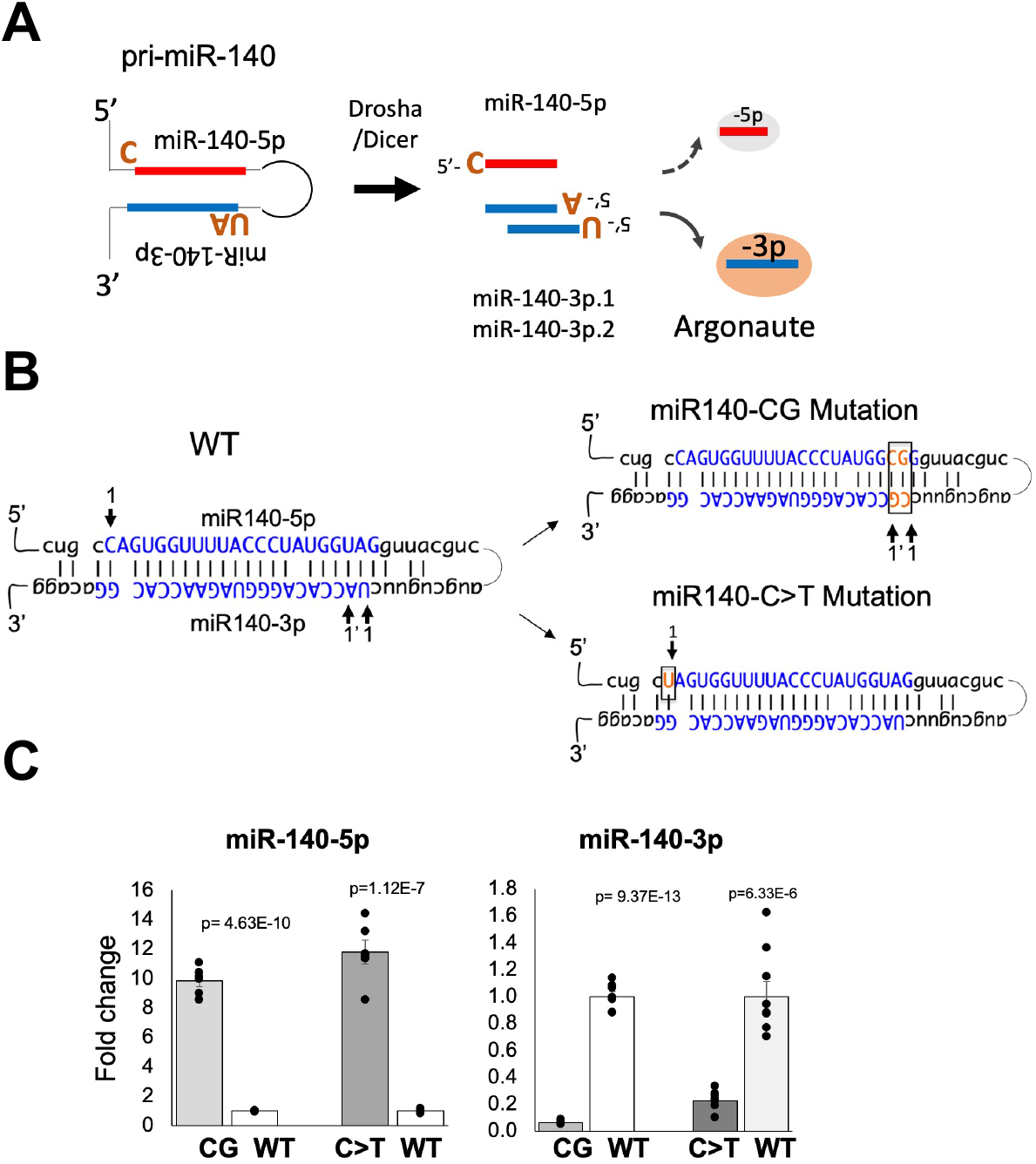
Method and validation of stoichiometric reversal in miR-140 miRNAs **A.** Schematic of miR-140-3p and -5p miRNA asymmetry by Argonaute (Ago) loading. The schematic shows that the three miR-140 miRNAs, miR-140-5p, miR-140-3p.1, and miR-140-3p.2, are derived from Drosha cleavage of pri-miR-140 and subsequent cleavage of the loop region of the pre-miR-140 by Dicer. Note that two miR-140-3p miRNAs are produced (miR-140-3p.1 and -3p.2) from the - 3p arm because Dicer cleavage occurs at two sites. The three miRNAs are asymmetrically loaded to Ago mostly dependent on the nature of the 5’ ends. Due to the preferential loading of miRNA strands with the 5’ nucleotides U or A, miR-140-3p miRNAs are more efficiently loaded to Ago than miR-140-5p that starts with C, a non-preferred nucleotide for Ago loading. **B.** Strategy to change the stoichiometry of miR-140-3p and -5p miRNAs. Blue bases indicate the mature miRNA sequences. Arrows indicate the first nucleotides of mature miRNAs. Base pairs in orange indicate where the mutations were made in each mouse model. The mutation from C to U of the 5’ nucleotide of miR-140-5p was designed to increase preferential Ago-loading of miR-140-5p in comparison to miR-140-3p miRNAs; this model was designated as miR-140-C>T. The second model was created by altering the 5’ nucleotides of the miR-140-3p strand to C and G from U and A, respectively, which are non-preferential for Ago-loading; this model was designated as miR-140-CG. Note that the 3’ end of miR-140-5p had to be altered in the miR-140-CG model to maintain the hairpin loop structure of the pri-miR-140 allowing for normal processing by Drosha and Dicer and to provide thermodynamic stability of the 5’ end of miR-140-3p of the miR-140-5p/-3p duplex upon cleavage. **C.** Relative expression of miR-140-3p and -5p in primary rib chondrocytes of miR-140-C>T, miR-140-CG, and respective WT littermates. miR-140-CG and miR-140-C>T mice show significantly increased miR-140-5p and significantly diminished miR-140-3p expression. Mean ± SEM with individual data is shown. N= 5-8 from 3 biological replicates.

To assess the effects of the miR-140-CG and miR-140-C>T mutations on the abundance of miR-140-5p and -3p, primary chondrocytes were isolated from P10 mice ribs. RNA was isolated and purified from these cells. As expected, real time-qPCR revealed a significant decrease in miR-140-3p expression and a significant increase in miR-140-5p expression in both the miR-140-CG and -C>T models (Fig. 1C).

### Reversing the stoichiometry of miR140-3p and miR140-5p leads to a mild delay in chondrocyte maturation with a delayed secondary ossification center formation

While stoichiometric inversion in these models was confirmed, the homozygous mice showed no overt gross abnormalities in growth (data not shown). However, histological assessment of the hindlimbs of these mice at postnatal day 10 revealed a consistent and mild delay in epiphyseal development in both models (Supplemental Fig. 1-2). The proximal tibias showed that in the mice homozygous for either miR140-CG or miR140-C>T a modest delay in chondrocyte maturation with delayed secondary ossification center development was presented (Fig. 2).

**Figure 2.**
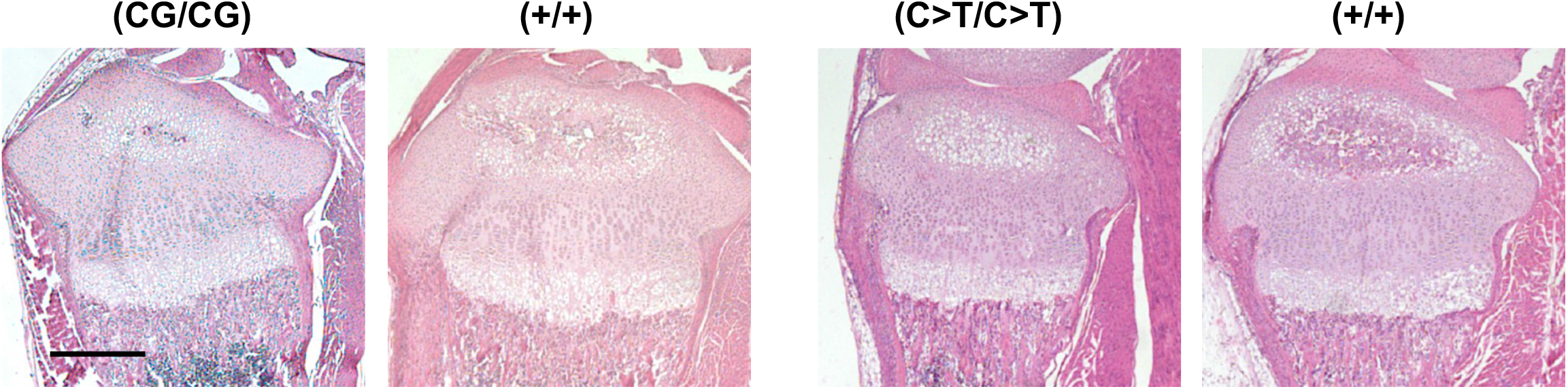
Histology of the proximal tibia of miR140-CG and −C>T models. Representative images of Hematoxylin and Eosin-stained proximal tibias of miR140-CG mice and miR140-C>T mice and control littermates at postnatal day 10. miR140-CG and −C>T tibias present a modest delay in the maturation of chondrocytes with delayed development of the secondary ossification center. Scale bar represents 500 um.

### The miR-140-C>T and -CG mutations enhance the phenotypic changes caused by the gain-of-function mutation of miR-140-5p

To further test -C>T and -CG mutations increase of miR-140-5p expression without changing the -5p function, we tested whether -C>T and -CG mutations enhanced the phenotypic changes of a mouse model for spondyloepiphyseal dysplasia (SED) MIR140 type. We previously demonstrated that a single nucleotide substitution (A to G; designated as miR-140-G mutation) of the first nucleotide of the seed sequence, *i.e*. the second nucleotide, of miR-140-5p caused a novel skeletal dysplasia and showed that the equivalent mutation in mice causes a delay in chondrocytes maturation in the epiphysis due to an expansion of resting zone chondrocytes (Grigelioniene et al. 2019) (Fig. 3A). This phenotype is caused by a gain-of-function of the mutant miR-140-5p in a dose dependent manner, but not by the loss of the function of the wildtype miR-140-5p. We hypothesized that the addition of the -C>T and -CG mutations would produce a skeletal phenotype similar to the G mutant with increased severity due to the increased expression of mutant miR-140-5p by the addition of -C>T and -CG mutations. To test this hypothesis, we changed the first two nucleotides of the miR-140-5p from C and A to U and G with the added -CG mutation; this mutation was designated as the miR-140-UGCG model (Fig. 3A). Mice heterozygous for the UGCG mutation showed a delay in secondary ossification center formation and an expansion of the resting zone of the growth plate similar to the G-mutation phenotype with greater severity (Fig. 3B, C). In addition, the UGCG homozygotes showed a more severe skeletal growth defect (Supplemental Fig. 3), unlike the relatively mild dwarfism and craniofacial deformity in homozygous G mutants (Grigelioniene et al. 2019). These data provide evidence supporting that the -C>T and -CG mutations do not alter the function of the mutant miR-140-5p, but rather increase its expression and the phenotypic severity of the gain-of-function mouse model.

**Figure 3.**
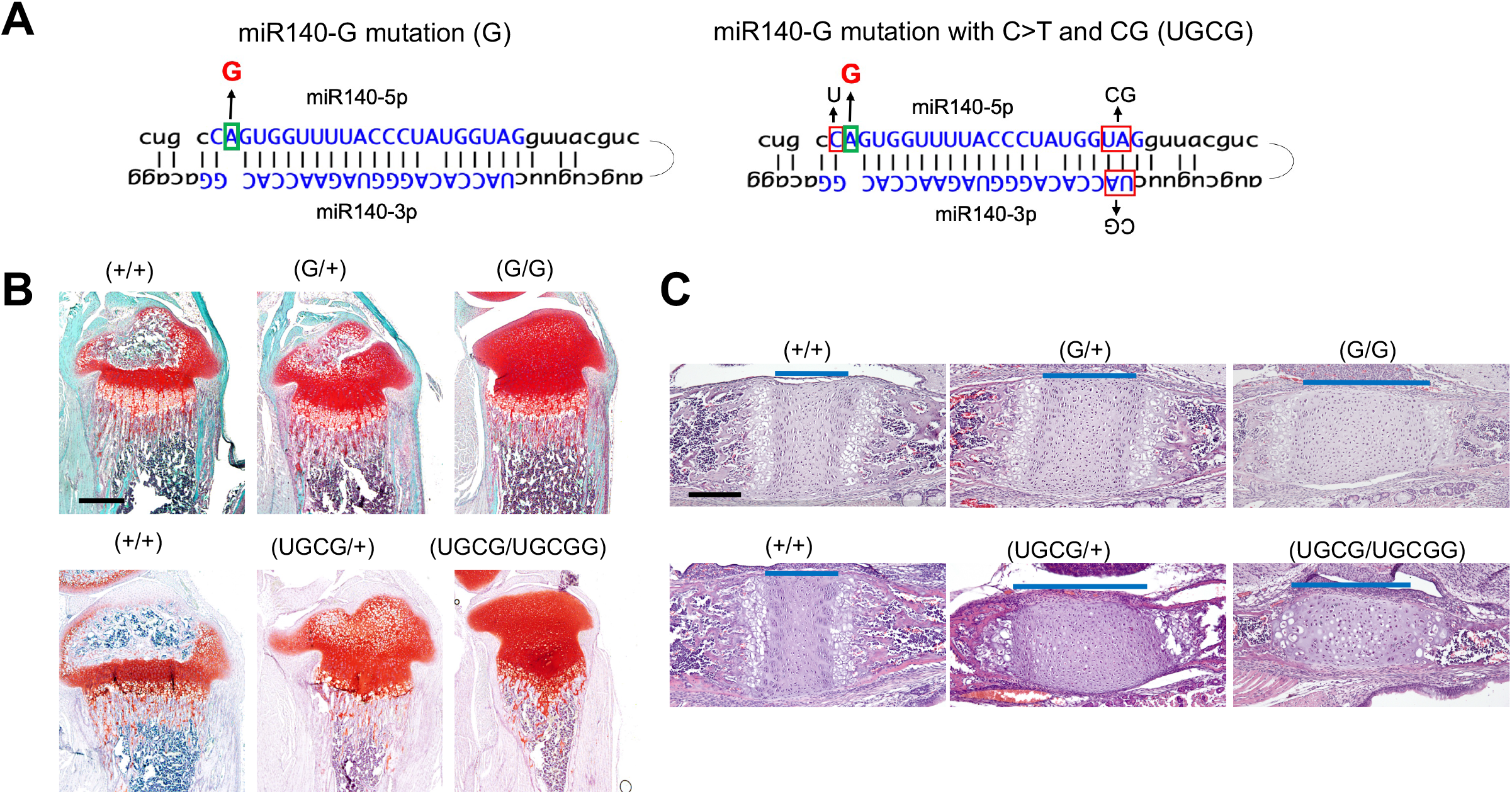
A gain-of-mutation in the C>T and CG context enhances the phenotype. **A.** A gain-of-function mutation at the second nucleotide of miR-140-5p (A>G) (G) was combined with C>T and CG mutation (UGCG). **B, C.** Safranin O-stained tibial sections of P14 mice (B) and hematoxylin and eosin-stained spheno-occipital growth plates in the basal skull of P7 mice (C) with indicated genotypes. The A>G mutation delays chondrocyte differentiation and epiphyseal development (B) and causes expansion of proliferating chondrocytes (blue lines in C). Additional UGCG mutations cause greater changes in the heterozygous state. Note the extent of expansion of the spheno-occipital growth plate of (UGCG/+) mice is greater than that of (G/+) mice and comparable to that of (UGCG/+) mice. Scale bars: A. 500 um, B. 200um

### RNA-sequencing analysis of miR-140-CG and miR-140-C>T primary chondrocytes reveals miR-140 targets

The effects of stochiometric alteration on gene expression was assessed by RNA-sequencing analysis using primary rib chondrocyte RNA from miR140-CG and miR140-C>T with respective controls (GSE162266). Additionally, the RNA-sequencing analysis of these two models was compared with a previous miR-140-knockout (KO) RNA-seq analysis (GSE98309). The stoichiometric change in these models is expected to increase and decrease the expression of miR-140-3p or miR-140-5p target genes, respectively. In order to test this hypothesis, we analyzed expression of predicted targets of miR-140-5p, miR-140-3p.1, and miR-140-3p.2. The target genes were predicted by TargetScan, and the target genes with 8-mer conserved binding sites of miR-140 miRNAs were used for analysis (Agarwal et al. 2015). A cumulative distribution fraction was drawn based on the expression changes of the target genes in these mice (Fig. 4A). As expected, the miR-140-5p target genes were significantly suppressed in both the miR-140-CG and -C>T models (Fig.4A). In contrast, both models present modest upregulation of miR-140-3p target genes (Fig. 4A). These results suggest that miR-140-3p miRNAs have a relatively weak regulatory effect on target genes, whereas miR-140-5p miRNA has a relatively greater regulatory effect, as suggested by our previous work on the miR-140-KO mouse analysis (Grigelioniene et al. 2019).

**Figure 4.**
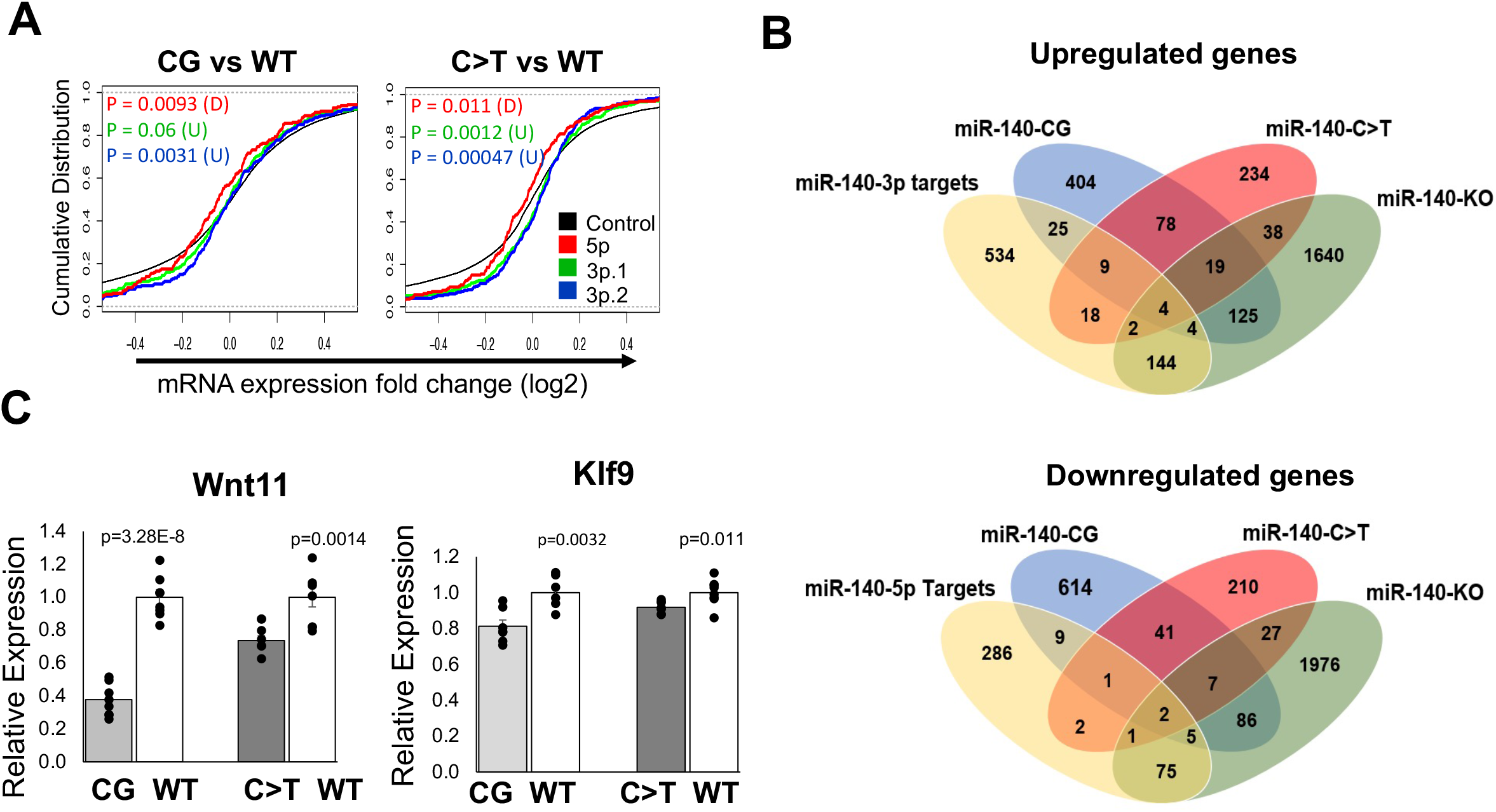
RNA-sequencing analysis of the miR-140-CG and −C>T homozygous chondrocytes reveals targets of miR-140 miRNA regulation. **A.** Cumulative distribution fraction analysis of miR-140 target gene expression in miR-140-CG and miR-140-C>T compared to control gene sets. Cumulative distributions of predicted miR-140-target genes with conserved 8mer-seed binding sites for miR-140-5p, -3p.1, and 3p.2 from miR-140-CG and respective WT chondrocytes and miR-140-C>T and respective WT chondrocytes were plotted based on their expression changes. The black lines indicate distribution of entire genes in these comparisons. The miR-140-5p target gene distribution curve is shifted to the left of the black line, indicating that the expression levels of predicted miR-140-5p target is generally suppressed in -CG or -C>T models. Likewise, miR-140-3p target gene distribution is modestly shifted to the right, demonstrating upregulation of miR-140-3p target genes in mutant chondrocytes. **B.** Two Venn diagrams were created to analyze the overlaps among the deregulated genes in miR-140-CG and miR-140-C>T models, the previously reported miR-140-KO model, and predicted miR-140-5p or miR-140-3p (−3p.1 and -3p.2) targets. Genes whose expression is altered by more than 20.5% were counted. miR-140 targets with conserved 8-mer seed binding sites were predicted by using TargetScan (v. 7.2). **C.** Relative expression of miR-140-5p targets, *Wnt11* and *Klf9*, in −CG and −C>T models Mean ± SEM, n=6 from 3 biological replicates.

The regulatory effects of miRNAs are estimated to be relatively modest, usually less than 50% and often less than 20% (Baek et al. 2008; Selbach et al. 2008). For this reason, we analyzed genes of which expression was altered by 20% or more in homozygous -CG or -C>T mutant mice and in miR-140-knockout (KO) mice. We found that 404, 234, and 1640 genes were upregulated in miR-140-CG, miR-140-C>T, and miR-140-KO mice, respectively. In addition, we revealed 614, 210, 1976 genes were downregulated in miR-140-CG, miR-140-C>T, and miR-140-KO mice, respectively (Fig. 4B). Additionally, TargetScan (v. 7.2) was used to predict the potential targets of miR-140 miRNAs by analyzing conserved 8mer sites and cumulative weighted score ++ of the miR-140 miRNAs (Agarwal et al. 2015). The overlap of these groups of genes are summarized in two Venn diagrams (Fig. 4B; Supplemental Table 1.). Among the miR-140-5p predicted targets, two genes, *Wnt11* and *Klf9*, were downregulated in -C>T and -CG mutants and upregulated in miR-140-KO mice. These genes are likely directly suppressed by miR-140-5p, and their expression is dependent on endogenous miR-140-5p. Likewise, among the predicted miR-140-3p miRNA targets, four genes of which expression was upregulated in all three mutants, -C>T, -CG, and -KO mice, were identified. These are likely direct miR-140-3p targets of which expression is dependent on miR-140-3p miRNAs. To confirm the predicted miR-140-5p targets, qRT-PCR was performed, which revealed the downregulation of *Wnt11* and *Klf9* expression in -C>T and -CG mutant chondrocytes (Fig. 4C).

### The role of miR-140-5p regulation on Wnt11 gene expression

Based on the mouse phenotypes where *Mir140* loss accelerates chondrocyte differentiation and -C>T and -CG mutants showing delays in chondrocyte differentiation in the secondary ossification center, we hypothesized that the loss of miR-140-5p-dependent regulation on the identified target genes might promote the secondary ossification center development and chondrocyte maturation. Since *Wnt11* was downregulated to a greater extent than *Klf9* in -C>T and -CG mutant chondrocytes and *Wnt11* contains only one predicted miR-140-5p binding site the 3’-UTR, we investigated the consequence of the disruption of the miR-140-5p binding site in mice. We disrupted this binding site by CRISPR/Cas9 genome editing in mice in two ways (Fig. 5A; Supplemental Fig. 4). One model, Wnt11-Del, was created by deleting a 14bp-long sequence including the miR-140-5p binding site using template-free genome editing (Fig. 5A; Supplemental Fig. 4). The second model, Wnt11-LoxP, was generated by inserting a loxP sequence ablating the miR-140-5p binding site (Fig. 5A; Supplemental Fig. 4). Primary chondrocytes were isolated from homozygous mice of both models, and RNA was subsequently isolated from these cells. We confirmed that both the Wnt11-Del and Wnt11-LoxP have significantly increased *Wnt11* expression, demonstrating that the miR-140-5p binding site is an important cis-regulatory sequence (Fig. 5B).

**Figure 5).**
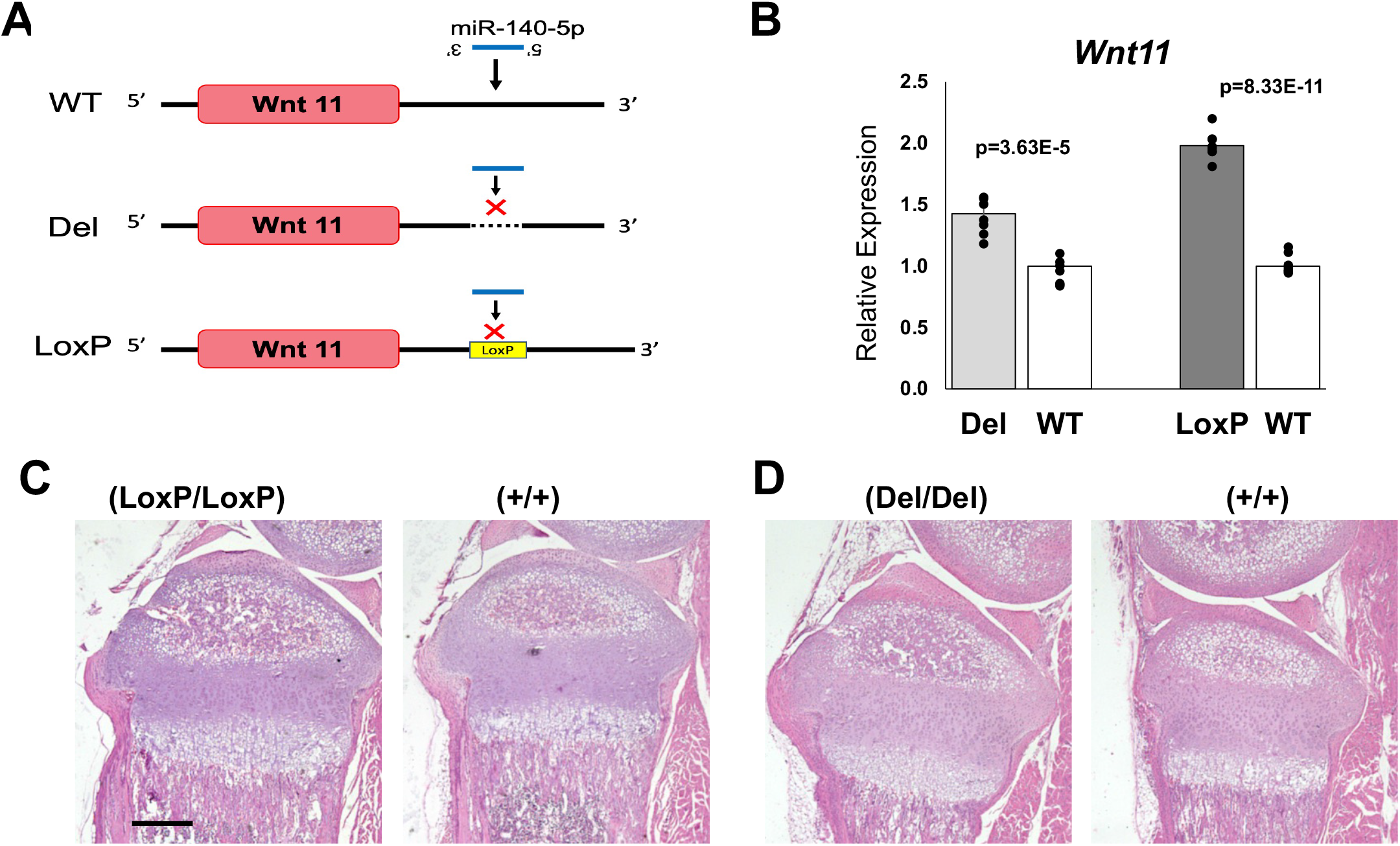
Generation and analysis of Wnt11-LoxP and Wnt11-Del mutant models. **A.** Schematic demonstrating the mutations made to the miR-140-5p binding site of the 3’-UTR of the Wnt11 gene. The Wnt11-Del model was created by deleting the 14 -bp sequence including the miR-140-5p binding site. The Wnt11-LoxP mutation was created by inserting a loxP sequence in the 3’-UTR ablating the miR-140-5p binding site. Both models were designed to diminish miR-140-5p binding. **B.** Relative expression of *Wnt11* in rib chondrocytes of homozygous Wnt11-LoxP and Wnt11-Del models. *Wnt11* expression is significantly increase in both models. Mean ± SEM with individual data is shown. N=6-8 from 3 biological replicates **C., D.** Representative images of hematoxylin and eosin-stained proximal tibias of Wnt11-LoxP (C), Wnt11-Del (D) and respective littermate controls. Both models had a modest acceleration of secondary ossification development (C-D). Scale Bar represents 500 um.

To assess the phenotypic outcome of these two models, sagittal sections of P10 mice hindlimbs were taken at 6 um to analyze the histology of these mice. Both the Wnt11-LoxP model and Wnt11-Del model showed a mild, but consistent, acceleration in the secondary ossification development compared to respective controls (Fig. 5C-D; Supplemental Fig. 5-6). It is important to note that a mild acceleration of endochondral bone development was found in neonatal *Mir140*-null mice, suggesting that *Wnt11* de-repression is responsible for this particular phenotype (Nakamura et al. 2011). Nevertheless, the overall skeletal phenotype is modest compared with *Mir140*-null mice, indicating that other physiologically important targets of miR-140-5p exist.

## Discussion

Here, we demonstrate that the miRNA-5p and -3p stoichiometry can be altered in mice by manipulating the 5’ nucleotides of miRNA arms. We have found that mice with these stoichiometrically reversed mutations show a delay in epiphyseal development. Additionally, through molecular analysis of these models, we identified target genes of miR-140-5p, including *Wnt11*. The removal of the miR-140-5p binding site of the 3’ untranslated region of *Wnt11* demonstrated the direct regulation of miR-140-5p on *Wnt11* expression and its role in controlling epiphyseal development. These results demonstrate that the stoichiometric inversion of miRNA - 5p and -3p is a useful approach not only for investigating the physiological roles of a given miRNA *in vivo*, but also for identifying target genes of miRNAs.

By changing the first 5’ nucleotide of miR-140 miRNAs, we reversed the stoichiometry of miR-140-5p and -3p and substantially increased the -5p abundance. The cumulative distribution fraction analysis also showed that the predicted miR-140-5p target genes were significantly suppressed. Yet, the physiological outcome of the stoichiometric reversal resulted in only a mild delay in the epiphyseal development, a far less severe phenotype when compared to the loss of miR-140-5p in *Mir140*-null mice (Nakamura et al. 2011). These results suggest that miR-140-5p is necessary for suppressing undesired genes for normal endochondral bone development, while overexpression of miR-140-5p has limited additional effects. Alternatively, it is possible that the simultaneous reduction of miR-140-3p expression might counteract the effect of miR-140-5p overexpression, although this is unlikely given the observation that the reduction of miR-140-3p had only a modest effect on the expression of predicted target genes. Previously reported findings revealed that the regulatory effect of miRNAs on each target is generally modest, usually less than 50%, and more often less than 20% (Baek et al. 2008; Selbach et al. 2008). In addition, the notion that miRNAs serve as a “genetic buffer” to suppress aberrant expression of undesired genes to ensure the robustness of cellular phenotype also predicts that miRNAs should not have very strong dose-dependent effects (Cassidy et al. 2013; Posada and Carthew, 2014). Consequently, the reported findings in combination with our results suggest that miR-140-5p reduces undesired genetic expression effects without many erroneous outcomes due to high concentration of miR-140-5p.

Although a few hundreds of genes are predicted as miR-140 targets by a prediction algorithm, the RNA-sequencing analysis of the miR-140-CG, -C>T, and -KO models identified only a limited number of target genes. Computational predictions do not address the cell-specific conditions, including the relative abundance of target transcripts and the miRNA of interest, and therefore, physiologically meaningful interactions between a miRNA and its targets need to be determined experimentally. Nevertheless, we identified only a limited number of targets genes for miR-140-5p and -3p. This result may be due to the sample variation outweighing the relatively modest effect on gene expression caused by the miR-140 manipulation thus reducing the sensitivity of the analysis. Although these limitations exist, the *Wnt11* models provide evidence that this approach can identify high confidence and physiologically meaningful targets of a miRNA. Additionally, it is important to note that a recent study analyzed the overexpression of miR-140-3p miRNAs in which miR-140-3p targets using human articular cartilage were identified, including *Rap1b, Aldh1a3, Cap1, Rhoc, Tnfsf12, Pkp1, Nid1, Chst14*, and *At1b2* (Woods et al. 2020). These identified miR-140-3p targets are consistent with those found in the analysis of our models, providing positive evidence for the validity of our approach. However, due to the cumulative distribution fraction presenting a relatively weak regulation of miR-140-3p miRNAs and suggesting a greater regulation for miR-140-5p targets, we focused on miR-140-5p targets, as we hypothesized these targets would present greater regulatory effects.

The *Wnt11* models validated the regulatory role of the miR-140-5p binding site in the 3’ UTR of the Wnt11 gene and suggested that *Wnt11* de-repression play a causal role in part of the phenotype of *Mir140*-null mice. *Wnt11* is a non-canonical Wnt-ligand and is expressed in pre-hypertrophic chondrocytes (Church et al. 2002; Witte et al. 2009; Kinsley et al. 2015). *Wnt11* overexpression generally stimulates chondrocyte marker expression in primary chondrocytes and mesenchymal progenitor cells (Bergwitz et al. 2001; Ryu and Chun, 2006; Liu et al. 2014), although in retroviral expression of Wnt11 in developing chick limbs shows a limited effect on chondrocyte differentiation (Church et al. 2002). It is, therefore, possible that the Wnt11 upregulation promotes the process of endochondral bone development in these models.

Although Wnt11 expression was upregulated in both the Wnt11-Del and -LoxP models, we found that the relative expression of *Wnt11* in the -LoxP model was greater than the Wnt11-Del model. In addition, the Wnt11-LoxP model showed acceleration of epiphyseal development more consistently than the Wnt11-Del model. Since both cases, the seed binding site is disrupted, we do not know the precise reason for this difference. It is possible that these minor genetic modifications might have created a new regulatory element for trans-acting regulators factors, such as RNA-binding proteins and other miRNAs, although we did not find known binding motifs or significant seed binding sites in the Del mutant sequence.

In summary, the reversing of the stoichiometry of miR-140-3p and -5p by manipulating the 5’ nucleotides of miR-140-3p and -5p miRNAs presents a useful approach to identifying targets of miRNA regulation and analyzing physiological roles of miRNAs in vivo. These results, in combination with further downstream analysis of the *Wnt11* models in which the models showed a modest acceleration in epiphyseal development when the miR-140-5p binding site is disrupted, suggests that miR-140-5p has only a modest regulatory effect on endochondral bone development. Additionally, the regulation of miR-140-5p on *Wnt11*, alone, only accounts for a portion of miR-140-5p’s overall regulation.

## Methods

The mouse experiments were approved by the institutional animal care & use committee (IACUC) of Massachusetts General Hospital.

### Animals

The genotyping of the miR-140-CG, miR-140-C>T, Super miR-140, and WNT11 Del and LoxP mice mutants was performed by using EconoTaq PLUS GREEN Master mix (Lucigen). The following primers were used for genotyping:

For detection of the Mir-140 CG mutant allele, Mir140-Loop-CG-F, CGGgttacgtcatgctgttcCG and Mir140-Loop-common-R ACCCAATAGACGCCTTAGCA. For Wt Mir140 allele, Mir140-Loop-Wt-F, GGtaGGTTACGTCA TGCTGTTCTA and Mir140-Loop common-R primer. For genotyping of miR-140 C>T genotyping, a 216-bp genomic PCR product using the primers, Mir140Crspr-F, TCTGTGTTCATCCCATCCTG and Mir140Crspr-R, ATGGAGTCCTTCGATGCAGA is digested with Bfa1 and Bsr1 (New England Biolabs) that specifically cut the C>T allele and WT allele, respectively. Wnt11 Del mutant genotyping was performed by PCR using the del mutant allele-specific primer set Wnt11m1-5’, GCCCACCACATGGGTTGTA and Wnt11 C-3’ AGAGGGATTGAAGTGAGCCA, and WT allele-specific Wnt11 C-5’ GGAACCACTAACTTGGGTTG and Wnt11 C-3’primer. The Wnt11 LoxP allele is detected by Wnt11 mut-3’ GCTATACGAAGTTATtccatgt and Wnt11 C-5’primer.

### Mouse genome editing

CRSPR/Cas9-mediated mouse genome editing was performed by the in vivo mouse genome editing method, i-GONAD, according to the procedure as described with some modifications (Ohtsuka M. Genome biol. 2018), using the CD-1 outbred strain (Charles River Laboratory). For miR-140-UGCG mutation, the founder was generated in a hybrid of CD-1 and C57B6 strains, and subsequently backcrossed to CD-1 to establish F1 lines. Briefly, at 0.7 day post coitum, the female was anesthetized by continuous inhalation of isoflurane. The back of the mouse was shaved, fur was removed with hair remover, and the skin was sterilized with alcohol preps. Through a 1-cm long skin incision, the skin and abdominal muscle layer were bluntly separated. A 0.5 cm incision was made to the muscle layer to access the abdominal cavity. The fat pad attached to the ovary was identified and pulled outside of the body to expose the oviduct. 1.5 ul of CRISPR solution containing 1 ug/ul of Cas9 (Sigma, or IDT) premixed with 10 mM two-piece guide RNA (crRNA and tracrRNA, gRNA) (IDT) and 1ug/ul of a single strand DNA repair template, was injected into the oviduct either by puncturing the oviduct or through the infundibulum. Then, in vivo electroporation was performed using the square pulse generator, BTX-820 with the electrode tweezers, CUY652P2.5X4 (Nepa Gene, Japan). The setting of the pulse generator was 50V, 5msec, 8 pulses, and the electrode gap was 0.5 cm. After suturing the muscle layer and skin, the pregnant mother was returned to the cage. Pups were genotyped for desired modifications, then mated with CD1 wild type mice to establish the F1 lines. Heterozygous F1 mice were intercrossed to generate homozygotes.

Single strand DNA repair templates were designed to have symmetric 31-35-nt long homologous sequence at both sides of the central region containing the desired mutations. The sequences of crRNAs and repair templates are as follows:

For Mir140 CG mutation, crRNA binding sequence for the guide RNA is 5’-ACGTCATGCTGTTCTACCAC-3’and the sequence of repair template is 5’-ctctctgtgtcctgcCAGTGGTTTTACCCTATGGCGGgttacgtcatgctgttcCGCCACAGGGTAGAACCACGGACagggtactggagc (Capital letters indicate miR-140-5p and miR-140-3p sequence, underlined letters are the introduced changes). For Mir140 C>T mutation, crRNA binding sequence for the guide RNA is 5’-CATAGGGTAAAACCACTGGC -3’and the sequence of repair template is 5’-ggctcccgccctgtgtgtctctctctgtgtcctgcTAGTGGTTTTACCCTATGGTAGgttacgtcatgctg-3’ (Capital letters indicate miR-140-5p and miR-140-3p sequence and underlined letter is the introduced change). For Mir140 UGCG mutation, the same guide RNA for the C>T mutation was used, and the sequence of the repair template is 5’-GTGGCTCCCGCCCTGTGTGTCTctctctgtgtcctgcTGGTGGTTTTACCCTATGGCGGgttacgtca tgctgtt-3’ (Capital letters indicate the mutant miR-140-5p sequence and underlined letters are introduced changes). This manipulation was performed using Mir140 CG mutant mice. For Wnt11 UTR mutations, the crRNA binding sequence is 5’-AAATTTACAACCCAAGTTAG-3’and the repair template sequence for LoxP insertion is 5’-TCTGGAATGTTCTTTGGGACCCTGTGCCCACCACATGGAATAACTTCGTATAGCATACATTATACGAAGTTATTGGGTTGTAAATTTTTATTTTCCTTCCCCTCTCCGTGGG-3’ (underlined letters indicate a LoxP sequence)

### Histological analyses

Mice were sacrificed at indicated ages, and the skull and tibias were isolated and fixed in 10% Formaldehyde/PBS. The tibias were decalcified in EDTA for ~2 weeks, and then the right hind limb was processed in paraffin. Sagittal sections of the right hind limb were taken at 6 um. Sections were stained with Hematoxylin and Eosin (H & E) or safranin-O according to the standard protocols.

### Primary Rib Chondrocyte Isolation and Cell Culture

Mice were sacrificed at P10. Primary rib chondrocytes were isolated from these mice as previously described with some modifications (Nakamura et al. 2011) Following overnight collagenase digestion, the cells were placed in 2 mL tubes, centrifuged, and the collagenase mixture was removed from the cells. Cells were resuspended in DMEM, 10% FBS, and 1% P/S. Cells were filtered through a 0.40 um strainer (Corning) into a 6 well plate (Corning). Cells were cultured overnight in DMEM-containing 10% FBS and antibiotics before harvesting.

### Real Time PCR

Relative expression levels of miR-140-3p and -5p were assessed by qRT-PCR using QuantStudio3. RNA was isolated from primary rib chondrocyte cells cultures by using the Directzol-RNA Mini Prep Kit (ZYMO). The purified RNA samples were transcribed to cDNA for miR140-3p and -5p using miRNA cDNA kit (QuantaBio) or TaqMan miRNA assay kit (ThermoFischer). qPCR primers for miR140-3p and -5p and U6 were used with the universal primer included in the QuantaBio kit.

Additional purified RNA was used for transcription to cDNA using the Verso cDNA synthesis kit using random hexamer (Thermo Scientific). RT-qPCR was performed using SYBR Green master mix (QuantaBio). The values were normalized using Actb.

Primer sequences used for RT-qPCR are as follows:

Wnt11-L 5’-CAGGATCCCAAGCCAATAAA-3’ and Wnt11-R 5’-GACAGGTAGCGGGTCTTGAG-3’; Klf9 -F 5’-GCAGTGAGCTCCACATTTCA-3’ and Klf9-R 5’-CGCTAGTGATGGCTGTCGTA, and Actb-L 5’-GCACTGTGTTGGCATAGAGG-3’ and Actb-R 5’-GTTCCGATGCCCTGAGGCTCTT-3’. For miRNA quantification was performed TaqMan microRNA assay kit (Thermofisher).

Wildtype and -C>T and -CG mutant miR-140-5p’s were quantified using the 5’ primer that binds to the common sequence, 5’-AGTGGTTTTACCCTATGG-3’. For Wt and CG mutant miR-140-3p quantification, the miR-140-3p-specific primer mixture, miR-140-3p

5’-YRCCACAGGGTAGAACCAC-3’ where Y and Rare equimolecular mixtures of C/T and A/G, respectively, was used. U6 was used as an internal control, U6-F 5’-CGCTTCGGCAGCACATATAC-3’.

### RNA-sequencing Analysis and Bioinformatics

Purified RNA (≥1ug) was submitted to BGI for RNA-seq analysis. The samples were sequenced using PE transcriptome sequencing on the BGIseq 400 (GSE162266). In addition to the miR140-CG and miR140-C>T RNA-seq analysis, miR140-KO RNA-seq analysis from a previous report was used to assess miR140 miRNA targets (GSE98309). Normalized FPKM (Fragments per Kilobase of transcript per Million) counts were at a threshold of 20.5% change was used. TargetScanMouse (v. 7.2) was used to predict miR-140-5p or miR-140-3p targets by taking into account the target site conservation and cumulative weighted context ++ score (Agarwal et al. 2015).

Cumulative distribution fractions were performed by using predicted miR-140 target genes with conserved 8mer miRNA building sites. The following gene set sizes were used: miR140-3p.1, n= 467, miR140-3p.2, n= 496, and miR140-5p, n= 434. P values were calculated by one-sided Kolmogorov-Smirnov (K-S) test for either: upregulation or downregulation. Default R studio’s empirical cumulative distribution fraction function was used to create the cumulative distribution fraction.

### Statistical Analysis

For all bar graphs, the data is expressed as the mean ± SEM. Statistical significance between two groups was determined by student t-test.

## Acknowledgments

We thank Dr. Masato Ohtsuka for practical advice on i-GONAD. We also thank Center for Skeletal Research (P30AR075042) for assistance in histological analysis. This study was supported by the NIH grant AR06645 (T.K.).

## Notes

### Competing Interest Statement

The authors have declared no competing interest.

