## Supplemental Materials for "Reversing the miRNA -5p/-3p stoichiometry reveals physiological roles and targets of miR-140 miRNAs"

**Supplemental Figures**

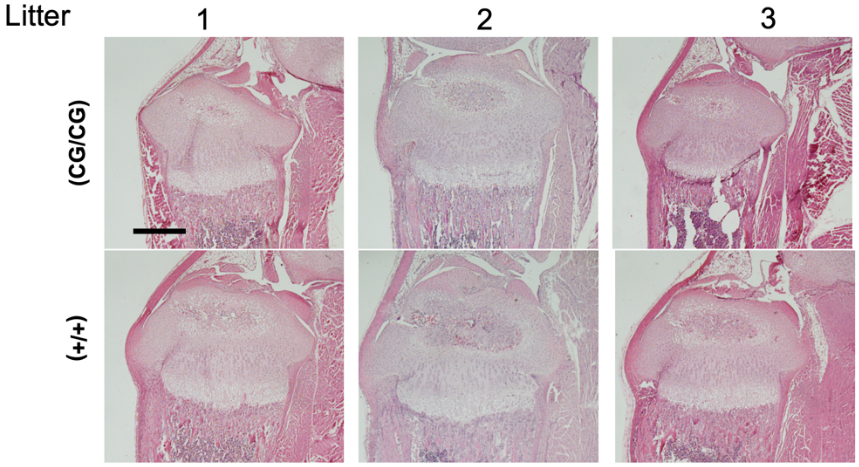


**Supplemental Figure 1.** Histology of hematoxylin and eosin-stained proximal tibias of miR-140-CG mice and littermate controls.   Scale bar presents 500 um.

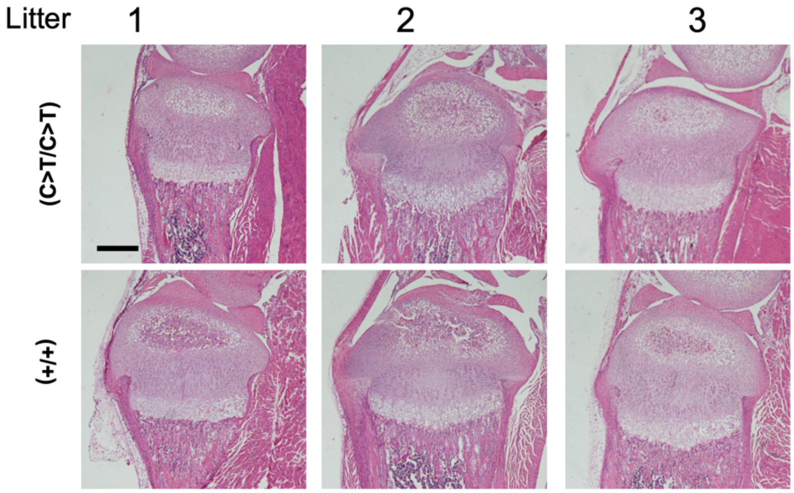


**Supplemental Figure 2.** Histology of hematoxylin and eosin-stained proximal tibia of miR-140-C>T and littermate controls. miR-140-C>T mutants were compared to littermate controls. Scale bar represents 500 um.

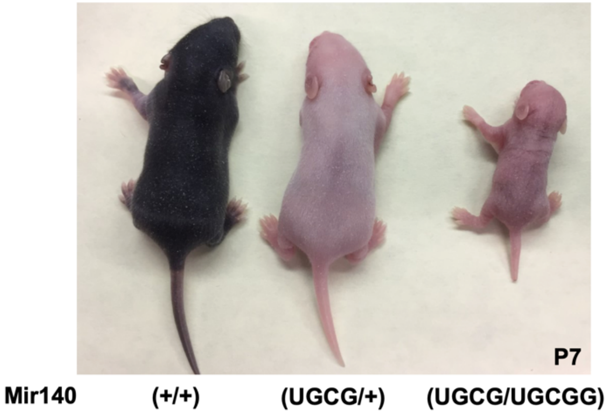


**Supplemental Figure 3.** Gross phenotype of miR-140-UGCG mutation. At P7, the homozygous Mir140(UGCG/UGCG) mutant mice show severely reduced body size unlike previously reported Mir140(G/G) mice. Mice are in a mixed background of CD1 and C57/B6.

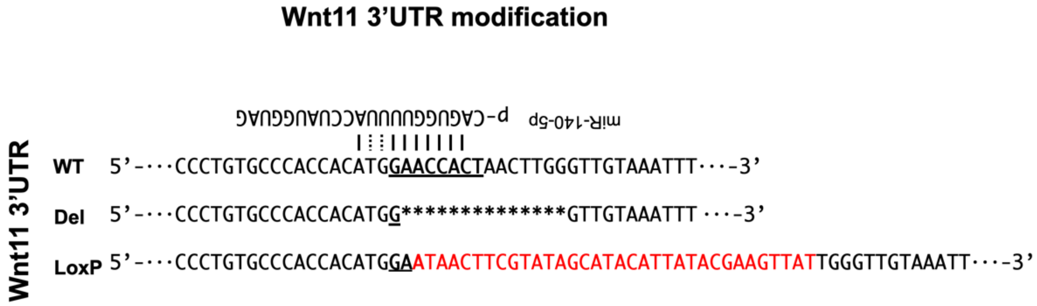


**Supplemental Figure 4.** Schematic of *Wnt11* 3'-UTR mutations. The WT miR-140-5p binding site sequence is shown with the corresponding miR-140-5p complementary sequence. Two models were made with different modifications. The Wnt11-Del allele has a 14-bp deletion including the predicted miR-40-5p binding site in the 3' UTR. The Wnt11-LoxP allele was created by adding a loxP sequence to ablating the miR-140-5p binding site.

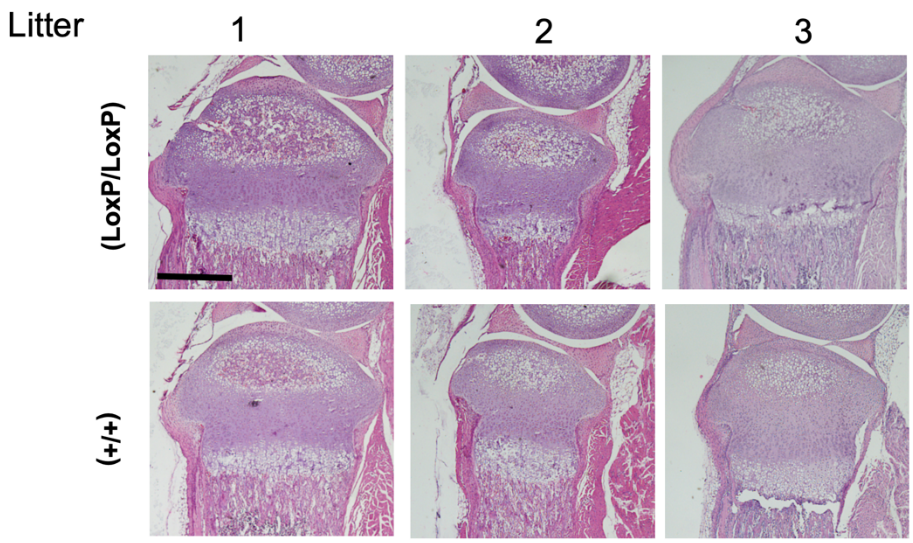


**Supplemental Figure 5.** Hematoxylin and eosin-stained proximal tibias of Wnt11-LoxP mice. Wnt11-LoxP mutants were compared to littermate controls showing a mild acceleration to the epiphyseal development of the proximal tibia consistently. Scale bar represents 500 um.

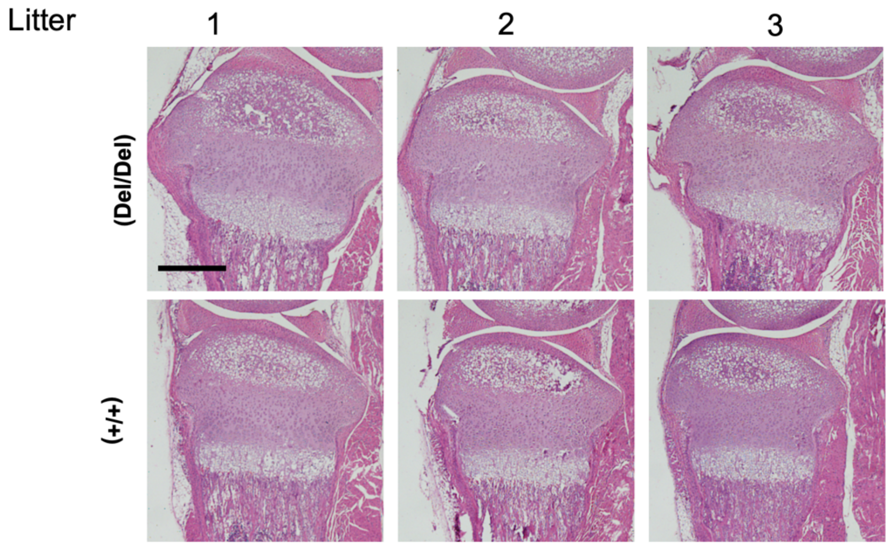


**Supplemental Figure 6.** Hematoxylin and eosin-stained Wnt11-Del mutant proximal tibias. The histology demonstrated that there is a modest acceleration in the epiphyseal development of the Wnt11-Del mutant mice in pairs 1 and 3 compared to littermate controls. Scale bar represents 500 um.

**Supplemental Table 1.** miR-140-3p and -5p miRNAs target lists compiled from TargetScan (v.7.2). The additional gene lists correspond to the Venn diagrams in Figure 4B.
